# Direct attenuation of Arabidopsis ERECTA signaling by a pair of U-box E3 ligases

**DOI:** 10.1101/2022.07.11.499617

**Authors:** Liangliang Chen, Alicia M. Cochran, Jessica M. Waite, Ken Shirasu, Shannon M. Bemis, Keiko U. Torii

**Author notes:** USDA-ARS Tree Fruit Research Laboratory, 1104 N Western Ave, Wenatchee, WA 98801, USA.

## Abstract

Plants sense a myriad of signals through cell-surface receptors to coordinate their development and environmental response. The Arabidopsis ERECTA receptor kinase regulates diverse developmental processes via perceiving multiple EPIDERMAL PATTERNING FACTOR (EPF)/EPF-LIKE peptide ligands. How the activated ERECTA protein is turned over is unknown. Here we identify two closely-related Plant U-box ubiquitin E3 ligases, PUB30 and PUB31, as key attenuators of ERECTA signaling for two distinct developmental processes: inflorescence/pedicel growth and stomatal development. Loss-of-function *pub30 pub31* mutant plants exhibit extreme inflorescence/pedicel elongation and reduced stomatal numbers owing to excessive ERECTA protein accumulation. Ligand-activation of ERECTA leads to phosphorylation of PUB30/31 by the co-receptor BRI1 ASSOCIATED KINASE1 (BAK1), which then promotes PUB30/31 to associate with and ubiquitinate ERECTA for eventual degradation. We further show that the PUB30 and PUB31 phosphorylation by BAK1 and their ubiquitination activities are critical for the proper *in vivo* developmental outputs. Our work highlights PUB30 and PUB31 as integral components of the ERECTA regulatory circuit that ensures optimal signaling strengths upon ligand activation, thereby enabling proper growth and development.

## Introduction

Development of multicellular organisms relies on coordinated cell proliferation and differentiation in response to external cues. Plants use a battery of membrane-bound cell surface receptors with intracellular kinase domain, collectively known as receptor-like kinases (RLKs), to sense and transduce external signals to adjust cellular responses^1^. Among them, those with extracellular leucine-rich repeat (LRR) domain, LRR-RLKs, comprise the largest subfamily in plants with more than 200 members in Arabidopsis^1^. The LRR-RLKs play critical roles in development, hormone perception, inter-kingdom communication and immunity. Those with known ligands will be referred as LRR-RKs hereafter^2,3^. Well-studied LRR-RKs include the brassinosteroid (BR) receptor BRASSINOSTEROID INSENSITIVE1 (BRI1)^4^ and the immune receptor FLAGELLIN SENSING2 (FLS2)^5,6^, among others^2,3^.

The Arabidopsis ERECTA LRR-RK regulates diverse aspects of plant development, including inflorescence architecture, stem and pedicel elongation, flower development, vascular differentiation, and stomatal patterning^7-12^. ERECTA perceives multiple peptide ligands, all belonging to EPIDERMAL PATTERNING FACTOR (EPF)/EPF-LIKE (EPFL) family^13^. Previous studies have identified both unique and shared components of ERECTA-signaling pathways. For example, among EPF/EPFL signaling peptides, EPFL6 (also known as CHALLAH) and EPFL4 from the stem endodermis are perceived by ERECTA and promote cell proliferation and stem/pedicel elongation for proper inflorescence architecture^14-16^. On the other hand, during stomatal development, EPF2 is primarily perceived by ERECTA to inhibit the stomatal-lineage entry divisions^17-19^. The receptor-like protein TOO MANY MOUTHS prevents signal interference between these EPF/EPFL-ERECTA mediated signaling to ensure proper stomatal patterning^15,16^. Upon ligand binding, ERECTA recruits universal co-receptor SOMATIC EMBRYOGENESIS RECEPTOR KINASES (SERKs), including SERK1, SERK3/BRI1-ASSOCIATED RECEPTOR KINASE1 (BAK1), and SERK4^20^. Immediate intracellular signaling components of ERECTA are shared for both inflorescence growth and stomatal development: those include receptor-like cytoplasmic kinases (RLCKs) BR-SIGNALING KINASE1/2 (BSK1/2) and a cascade of mitogen-activated protein kinases (MAPKs), YODA (YDA)-MKK4/5-MPK3/6^10,21-24^. SERKs/BAK and BSKs were originally identified as components of BR-activated BRI1 receptor complex^25-27^. The flagellin-activated FLS2 also form a complex with its co-receptor BAK1^28^ and the signal is then mediated by the MAPK cascades^29^.

After receptor activation, the strength of cellular signaling must be promptly downregulated to avoid excessive or untimely signal outputs. Thus, the mechanism of signal downregulation is an integral part of receptor signaling. Studies on FLS2 and BRI1 have highlighted the role of receptor ubiquitination (ubiquitylation) for signal attenuation: interestingly, both ligand-activated FLS2 and BRI1 are ubiquitinated by the two identical <u>P</u>lant <u>U</u>-<u>B</u>ox ubiquitin E3 ligases, PUB 12 and PUB13, albeit in a slightly different manner^30,31^. Whereas PUB12/13 target several additional RKs, ERECTA is not ubiquitinated by PUB12/13^31^. The U-box domain, which was originally identified as Ub Fusion Degradation 2 (UFD2) in yeast, mediates interaction with the ubiquitin-conjugating enzyme^32-35^. The first reported PUB protein is <u>A</u>RM <u>R</u>epeat <u>C</u>ontaining1 (ARC1), which interacts with the kinase domain of Brassica *S-locus* RKs^36^. The PUB proteins constitute a family of over 60 members in Arabidopsis, some of which are involved in a variety of environmental responses^37^. However, with the exception of a handful of members, their *in vivo* targets and functions remain unknown. Likewise, whether PUB proteins mediate the degradation of other LRR-RKs, or plant receptor kinases more broadly, remains an open question.

Here, we report two paralogous PUB proteins, PUB30 and PUB31, as key attenuators of ERECTA signal transduction pathways for both inflorescence/pedicel growth and stomatal development. The *pub30 pub31* double mutant plants exhibit characteristic inflorescence with extreme pedicel elongation and reduction in stomatal development. The *erecta* mutation is epistatic to *pub30 pub31*, indicating that PUB30 and PUB31 are redundantly required to downregulate ERECTA activity. We demonstrate that perception of EPF2 and EPFL6 peptides by ERECTA leads to the phosphorylation of PUB30/31 by BAK1 and stronger associations of PUB30/31 with ERECTA and BAK1. PUB30/31 directly ubiquitinate ERECTA for eventual degradation. Through site-directed mutagenesis, we further show that the PUB30 and PUB31 phosphorylation by BAK1 and their ubiquitin E3 ligase activities are critical for proper inflorescence/pedicel growth and stomatal development. Our work reveals the mode of actions and functions of a pair of PUB proteins in two ERECTA-mediated developmental processes and further suggests a broader view of how plant receptor kinases are attenuated upon signal activation.

## Results

### *PUB30/31* negatively regulate ERECTA-mediated inflorescence and pedicel growth

Loss-of-function *erecta* mutant plants exhibit characteristic compact inflorescence and short pedicels (Fig. 1a-c)^7,8,14,38,39^. We hypothesized that potential negative regulators of ERECTA may confer the opposite phenotype - i.e. extreme elongation of inflorescence and pedicels. With this in mind, we systematically surveyed the T-DNA insertion lines of *PUB* family genes. This led to the identification of *PUB30* and *PUB31* null mutant alleles (see Methods, Supplementary Fig. 1). Whereas single mutants of *pub30* and *pub31* do not show obvious growth phenotypes, the *pub30 pub31* double mutants produce elongated inflorescence with extremely long pedicels (Fig. 1a-c). Introduction of wild-type *PUB30* or *PUB31* coding sequences driven by their native promoters (*proPUB30::PUB30* and *proPUB31::PUB31*) into the *pub30 pub31* double mutant fully rescued the elongated pedicel phenotype (Supplementary Fig. 2a, b). These results indicate that *PUB30* and *PUB31* act redundantly to restrict the elongation of inflorescences and pedicels.

**Fig. 1.**
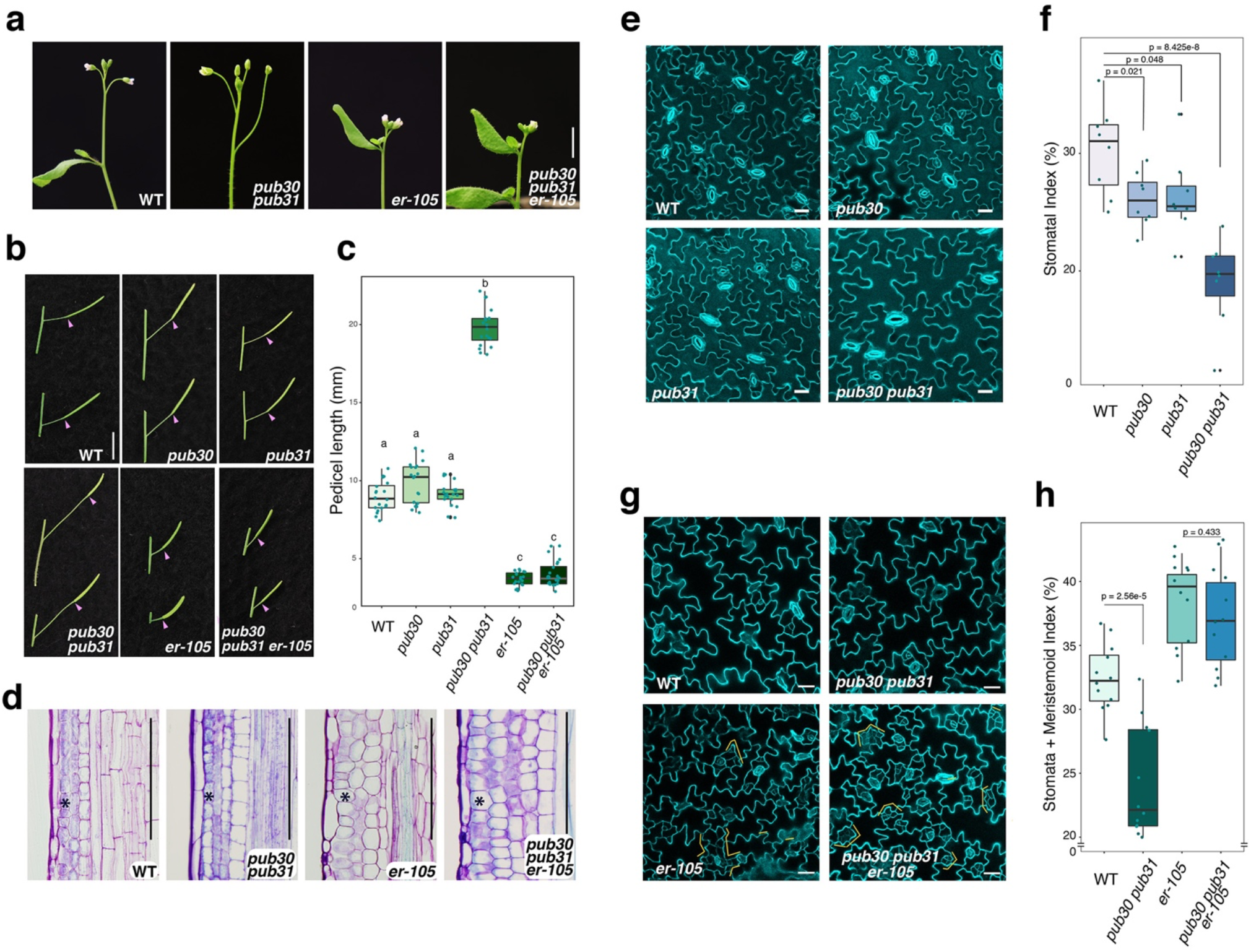
*PUB30/31* regulate inflorescence/pedicel growth and stomata development in the *ERECTA* pathway. (a) Representative inflorescence of wild type, *pub30 pub31, er-105*, and *pub30 pub31 er-105* plants. (b) Representative pedicels with fully-expanded siliques of wild type, *pub30, pub31, pub30 pub31, er-105*, and *pub30 pub31 er-105* plants. Scale bar: 1 cm (c) Morphometric analysis of pedicel length from each genotype. 6-wk-old mature pedicels (n = 20) were measured. One-way ANOVA followed by Tukey’s HSD test was performed for comparing all other genotypes and classify their phenotypes into three categories (a, b, and c). (d) Longitudinal sections of mature pedicels of wild type, *pub30 pub31, er-105*, and *pub30 pub31 er-105* plants. Asterisks, representative cortex cells in each genotype. Scale bar: 200 μm (e) Confocal microscopy of 10-d-old abaxial cotyledon epidermis of wild type, *pub30, pub31*, and *pub30 pub31*. Scale bar: 25μm (f) Quantitative analysis. Stomatal index (SI) of the cotyledon abaxial epidermis from 10-d-old seedlings of respective genotypes. Welch’s two sample t-test was performed for the pairwise comparisons with the wild type. p values are indicated in the graph. (g) Confocal microscopy of 6-d-old abaxial cotyledon epidermis of wild type, *pub30 pub31, er-105*, and *pub30 pub31 er-105*. Scale bar: 25 μm. (h) Quantitative analysis. Stomata + Meristemoid index of the cotyledon abaxial epidermis from 6-d-old seedlings of respective genotypes. Welch’s two sample t-test was performed for the pairwise comparisons of wild type vs. *pub30 pub31* as well as *er-105* vs. *pub30 pub31 er-105*. p values are indicated in the graph.

To address the genetic relationships of *PUB30/31* with *ERECTA*, we next generated the triple mutant of *pub30 pub31 er-105* (*erecta* null allele). The pedicel length of the triple mutant phenocopied that of *er-105* (Fig. 1a-c), indicating that the *erecta* mutation is epistatic to *pub30 pub31*. To further examine the underlying cellular basis of the *pub30 pub31* defects and its relationship with *erecta*, we analyzed the longitudinal sections of mature pedicels (Fig. 1d). It has been shown that short *erecta* pedicels accompany reduced cell proliferation and compensatory cell growth in the cortex layer ^14,38^. In contrast to *erecta*, cortex cells in the *pub30 pub31* pedicels are small and highly organized (Fig. 1d, asterisks), indicating that the extremely elongated pedicel phenotype of *pub30 pub31* is due to excessive cell proliferation, but not cell expansion. The *pub30 pub31 er-105* pedicels exhibit large, expanded cortex cells, thus phenocopying *er-105*. Thus, the *erecta* mutation is epistatic to not only the overall pedicel length but also the underlying cortex cell proliferation phenotype of *pub30 pub31*. Combined, our results suggest that *PUB30* and *PUB31* function as negative regulators of ERECTA-mediated inflorescence and pedicel growth.

### *PUB30/31* negatively regulate ERECTA-mediated inhibition of stomatal development

It is well known that ERECTA-family LRR-RKs enforce stomatal patterning^9^. Among the three members, ERECTA plays a major role in restricting the initiation of stomatal-cell lineages^9,19^. To dissect the genetic relationship between *ERECTA* and *PUB30/31* in stomatal development, we first analyzed the cotyledon epidermal phenotype (Fig. 1e, f). The *pub30* and *pub31* single mutants showed slightly reduced stomatal index (SI = number of stomata/(number of stomata + non-stomatal epidermal cells) x 100) compared with wildtype. The SI was significantly reduced in the *pub30 pub31* double mutant (Fig. 1e, f). Again, transgenic *pub30 pub31* plants expressing *proPUB30::PUB30* and *proPUB31:PUB31* fully rescued the stomatal phenotype of *pub30 pub31* (Supplementary Fig. 2c, d), indicating that *PUB30* and *PUB31* redundantly promote stomatal development.

We further characterized the stomatal phenotypes of *erecta, pub30 pub31*, and *pub30 pub31 erecta*. Consistent with the epistatic effect of *erecta* over *pub30 pub31* with respect to inflorescence and pedicel growth (Fig. 1a-c), *erecta* is epistatic to *pub30 pub31* on stomatal development: *er-105* confers increased numbers of small stomatal lineage cells (Fig. 1g, orange brackets) thus vastly elevating the Stomatal + Meristemoid index (number of stomata + meristemoids)/(number of stomata + non-stomatal epidermal cells) x 100) (Fig. 1h) due to excessive asymmetric entry division events^9,19^. The *pub30 pub31 er-105* epidermis is phenotypically indistinguishable from the *er-105* epidermis (Fig. 1g, h). Thus, *PUB30/31* negatively regulate two distinct *ERECTA*-mediated developmental processes: inflorescence/pedicel elongation and stomatal-lineage development.

### Ligand perception promotes the physical interaction of ERECTA and PUB30/31

*ERECTA* functions in the same genetic pathway with *PUB30/31* to regulate pedicel growth and stomatal lineage development (Fig. 1). Just like previously-reported localization patterns of ERECTA-YFP^40^, the YFP-fused PUB30 and PUB31 expressed by their own native promoters (*proPUB30::PUB30-YFP* and *proPUB31::PUB31-YFP*) are detected in the developing cotyledon epidermis, marking the plasma membrane (Supplementary Fig. 3a). The similar expression and localization patterns of ERECTA and PUB30/31 imply their potential interaction. To test whether ERECTA directly interacts with PUB30/31, we first performed a yeast two-hybrid assay (Y2H). A truncated ERECTA protein with cytosolic domain (ERECTA_CD), which contains the JMD, kinase domain and the C-terminal tail, was fused to the DNA binding domain (BD) and used as a bait. PUB30/31 are predicted cytoplasmic proteins, which contain a U-box domain, an ARM repeats, and a linker domain in between (Supplementary Fig. 3b). Full-length PUB30 and PUB31 proteins were fused to the activation domain (AD). As shown in Fig. 2a, ERECTA_CD interacts with PUB30/31. Next, we confirmed the direct interaction of ERECTA_CD and PUB30/31 by *in vitro* pull-down assay using purified recombinant ERECTA_CD and full-length PUB30 and PUB31 proteins (Supplementary Fig. 3c). Finally, to quantitatively characterize the kinetics of protein-protein interactions between PUB30/31 and ERECTA_CD, we performed biolayer interferometry (BLI) assays (see Methods). PUB30 and PUB31 bind with ERECTA_CD at a micromolar affinity (Fig. 2b, c), which appears too high to be considered for specific interactions.

**Fig. 2.**
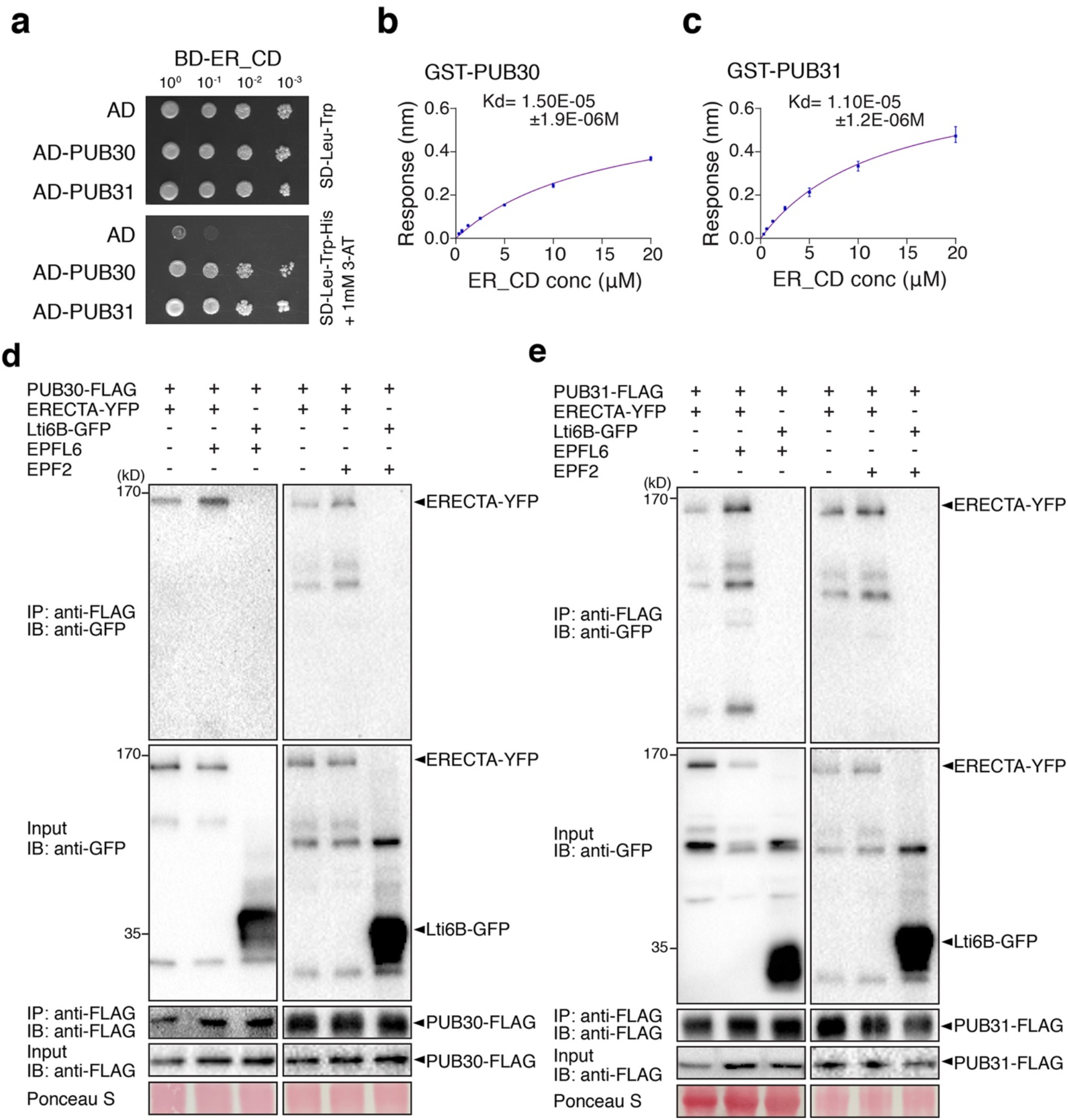
PUB30/31 directly interact with ERECTA. (a) PUB30 and PUB31 interact with the cytoplasmic domain of ERECTA (ER_CD) in yeast. ER_CD were used as bait. The activation domain (AD) alone, PUB30 and PUB31 were used as prey. Yeast were spotted in 10-fold serial dilutions on appropriate selection media. The experiment was repeated independently three times with similar results. (b) A quantitative analysis of interactions between PUB30 and ER_CD using BLI. *In vitro* binding response curves for recombinantly purified GST–PUB30 and MBP-ER_CD at seven different concentrations (312.5 nM, 625 nM, 1250 nM, 2500 nM, 5000 nM, 10000 nM, and 20000 nM) are shown. Kd values are indicated. Data are mean ± s.d., representative of two independent experiments. (c) A quantitative analysis of interactions between PUB31 and ER_CD using BLI. *In vitro* binding response curves for recombinantly purified GST–PUB31 and MBP-ER_CD at seven different concentrations (312.5 nM, 625 nM, 1250 nM, 2500 nM, 5000 nM, 10000 nM, and 20000 nM) are shown. Kd values are indicated. Data are mean ± s.d., representative of two independent experiments. (d) Both EPFL6 and EPF2 induce the association of PUB30 with ERECTA *in vivo*. After treatment with the ligands of ERECTA, EPFL6 and EPF2, proteins from *proPUB30::PUB30-FLAG*; *proERECTA:: ERECTA-YFP* and *proPUB30::PUB30-FLAG*; *Lti6B-GFP* plants were immunoprecipitated with anti-FLAG beads (IP), and the immunoblots (IB) were probed with anti-GFP and anti-FLAG antibodies, respectively. ERECTA-YFP was detected in the immunoprecipitated PUB30-FLAG complex. (e) Both EPFL6 and EPF2 induce the association of PUB31 with ERECTA *in vivo*. After treatment with the ligands of ERECTA, EPFL6 and EPF2, proteins from *proPUB31::PUB31-FLAG*; *proERECTA:: ERECTA-YFP* and *proPUB31::PUB31-FLAG*; *Lti6B-GFP* plants were immunoprecipitated with anti-FLAG beads (IP), and the immunoblots (IB) were probed with anti-GFP and anti-FLAG antibodies, respectively. ERECTA-YFP was detected in the immunoprecipitated PUB31-FLAG complex.

To examine the *in vivo* association of ERECTA with PUB30 and PUB31 in Arabidopsis, we further performed co-immunoprecipitation (Co-IP) analyses using transgenic plants carrying epitope tagged ERECTA (*proERECTA::ERECTA-YFP*) and PUB30/31 (*proPUB30::PUB30-FLAG* and *proPUB31::PUB31-FLAG*). Consistent with the *in vitro* BLI assays (Fig. 2b, c), ERECTA-YFP was barely detectable in the immunoprecipitated PUB30-FLAG or PUB31-FLAG complexes (Fig. 2d, e). It has been shown that EPF/EPFL ligand perception by ERECTA triggers the formation of an active receptor complex^20,41,42^. To test the hypothesis that receptor activation promotes the interaction of ERECTA and PUB30/31, we next treated the seedlings with EPFL6 and EPF2 peptides. Indeed, ERECTA strongly associates with PUB30 and PUB31 upon peptide stimulation (Fig. 2d, e). Combined, our results demonstrate that ERECTA physically interacts with PUB30 and PUB31, and their *in vivo* interactions are stimulated by corresponding peptide ligands regulating inflorescence elongation and stomatal development.

### PUB30 and PUB31 ubiquitinate ERECTA

As members of the PUB protein family, PUB30 and PUB31 possess sequence features of E3 ligases (Supplementary Fig. 4a). To determine whether PUB30/31 possess E3 ubiquitin ligase activity and whether ERECTA is their substrate, we first performed *in vitro* ubiquitination assays. Faint laddering bands of ERECTA (MBP-ERECTA_CD) were detected when co-incubated with PUB30/31 proteins (GST-PUB30 and GST-PUB31), E2 ubiquitin-conjugating enzyme (His-UBC8), and E1 ubiquitin activating enzyme (His-UBA1) (Supplementary Fig. 4b, c), indicating that PUB30/31 can ubiquitinate ERECTA *in vitro*.

To address the *in vivo* role of PUB30/31 in regulating ERECTA, we next compared the *in vivo* ubiquitination status of ERECTA in *erecta* null mutant, *er-105*, complemented with *proERECTA::ERECTA-FLAG* (hereafter referred to as ‘wild type’) and *erecta pub30 pub31* triple mutant complemented with *proERECTA::ERECTA-FLAG* (*i*.*e. ‘pub30 pub31’*) seedlings^43^ (Fig. 3a, b). The relative signal intensity ratio between ubiquitinated ERECTA detected by α-ubiquitin antibodies and immunoprecipitated ERECTA detected by anti-FLAG antibodies, indicates that, in the absence *of PUB3/31*, ERECTA is less ubiquitinated *in vivo* (Fig. 3a, b). Poly-ubiquitinated proteins can be destined for degradation via the 26S proteosome pathway^44-46^. We subsequently examined whether PUB30/31 regulate ERECTA stability *in vivo*. Higher accumulation of ERECTA proteins (ERECTA-FLAG) was detected in *pub30 pub31* mutant background compared to the wild type (Fig. 3c, d). In contrast, *ERECTA* transcript levels were not significantly different between wild-type and *pub30 pub31* seedlings (Supplementary Fig. 4d), indicating that the effects of *PUB30/31* on ERECTA accumulation is likely post-translational. Treatment with the 26S proteasome inhibitor MG132 resulted in the significant increase in ERECTA protein accumulation in the ‘wild-type’ seedlings (Fig. 3c), indicating that the 26S proteosome pathway regulates ERECTA protein stability.

**Fig. 3.**
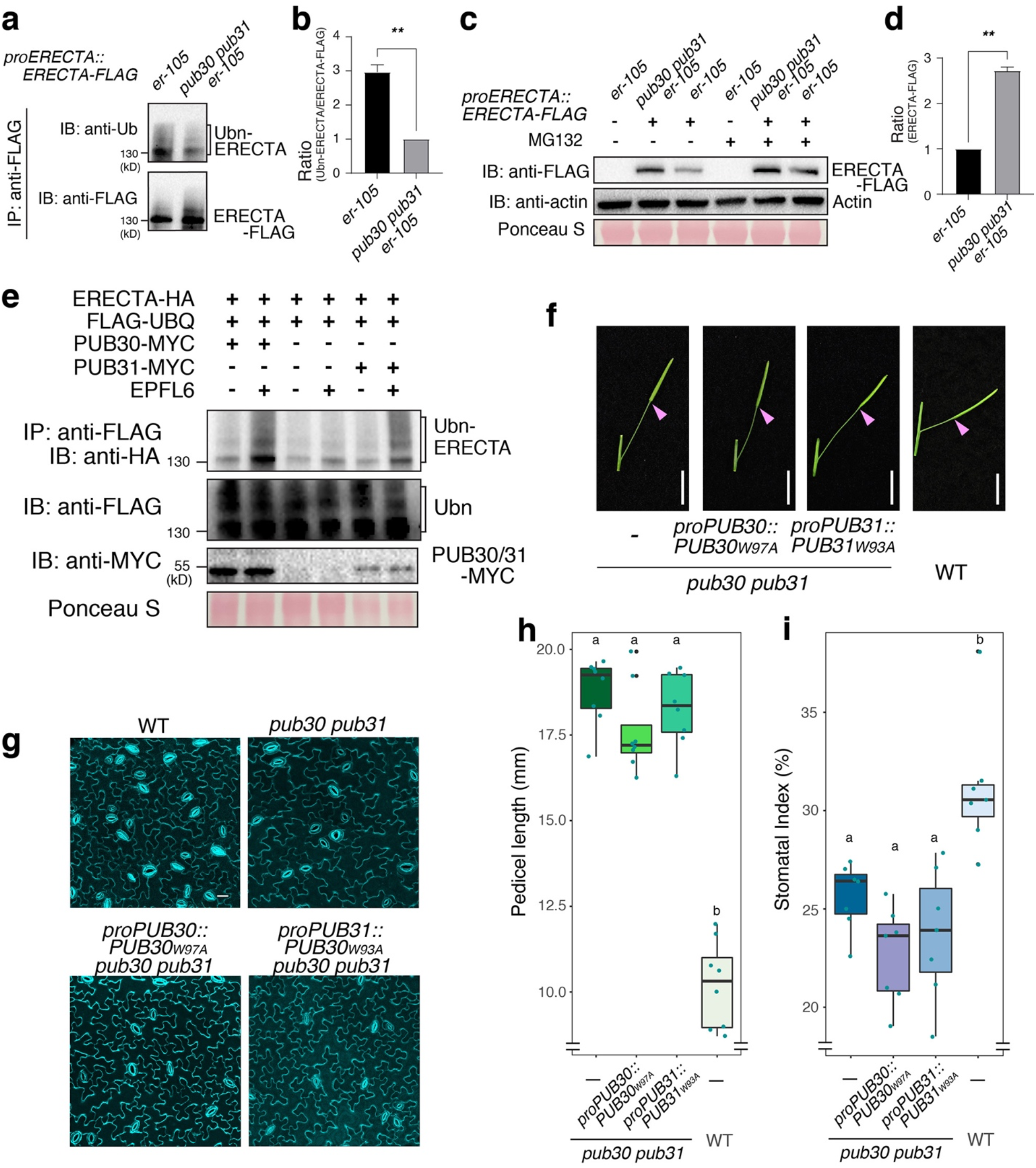
PUB30/31 ubiquitinate and regulate the protein abundance of ERECTA. (a) Reduced ERECTA ubiquitination in *pub30 pub31*. IP was performed using α-FLAG antibodies on solubilized microsomal fraction protein extracts from homozygous plants expressing ERECTA-FLAG in either wild-type or *pub30 pub31* background and the immunoblots (IB) were probed with anti-ubiquitin (P4D1) and anti-FLAG antibodies, respectively. (b) Quantitative analysis of ERECTA ubiquitination profiles. Error bars represent SD (n = 3). The asterisks indicate statistical significance by using Student’s t test (*P < 0.05). (c) ERECTA protein accumulation in wildtype, and *pub30 pub31*, in the absence and presence of the protease inhibitor MG132. Total membrane proteins were isolated from 7-d-old seedlings and probed by an α-FLAG antibody. The protein inputs were equilibrated using α-Actin antibodies. (d) Quantification of ERCTA abundance (ERECTA/Actin) (n = 3 biological replicates). The asterisks indicate statistical significance by using Students t test (*P < 0.05). (e) PUB30 or PUB31 mediates ERECTA ubiquitination *in vivo*. Arabidopsis protoplasts were cotransfected with HA-tagged ERECTA (ERECTA-HA), FLAG-tagged ubiquitin (FLAG-UBQ), and together with a control vector or MYC-tagged PUB30 or PUB31 (PUB30-MYC or PUB31-MYC) and incubated for 8 h followed by treatment with 5 μM EPFL6 for 1 h in the presence of 2 μM MG132. After immunoprecipitation using anti-FLAG beads, the ubiquitinated ERECTA was probed with α-HA antibody. The total ubiquitinated proteins were probed by an α-FLAG antibody and PUB30 or PUB31 proteins were probed by an α-MYC antibody. (f) Representative pedicels of mature siliques of *pub30 pub31, proPUB30::PUB30W97A*; *pub30 pub31, proPUB31::PUB31W93A*; *pub30 pub31*, and wild type plants. Scale bar: 1 cm (g) Confocal microscopy of 8-d-old abaxial cotyledon epidermis of *pub30 pub31, proPUB30::PUB30W97A*; *pub30 pub31, proPUB31::PUB31W93A*; *pub30 pub31*, and wild type plants. Scale bar: 25 μm (h) Morphometric analysis of pedicel length from each genotype. 6-wk-old mature pedicels (n = 15) were measured. One-way ANOVA followed by Tukey’s HSD test was performed for comparing all other genotypes and classify their phenotypes into two categories (a and b). (i) Quantitative analysis. Stomatal index (SI) of the cotyledon abaxial epidermis from 8-d-old seedlings of respective genotypes. One-way ANOVA followed by Tukey’s HSD test was performed for comparing all other genotypes and classify their phenotypes into two categories (a and b).

We further performed an *in vivo* ubiquitination assay using Arabidopsis protoplasts co-expressing epitope-tagged ERECTA (ERECTA-HA), PUB30 or PUB31 (PUB30-MYC or PUB31-MYC), and ubiquitin (FLAG-UBQ) (see Methods). Laddering bands with high-molecular-weight proteins are detected after immunoprecipitation (IP), indicative of the ubiquitination of ERECTA *in vivo* (Fig. 3e). Strikingly, application of EPFL6 peptide intensified the poly-ubiquitination of ERECTA by PUB30/31 (Fig. 3e), suggesting that PUB30/31 mediate the ligand-stimulated ERECTA ubiquitination.

Finally, to address whether the ubiquitination activity of PUB30/31 is essential for their function as regulators of ERECTA-mediated processes, we introduced amino-acid substitutions to PUB30/31 sequences that replace the conserved E2-binding tryptophan residue to alanine (PUB30_W97A_ or PUB31_W93A_) within their U-box motif (Supplementary Fig. 4a). *In vitro* autoubiquitination assays showed that these mutations (W97A in PUB30 or W93A in PUB31) diminished the ubiquitination activity of PUB30 and PUB31 (Supplementary Fig. 4e). Next, these E2-binding defective PUBs were expressed by their native promoters in the *pub30 pub31* mutant. Neither transgenic *proPUB30::PUB30*_*W97A*_ nor *proPUB31::PUB31*_*W93A*_ was able to rescue the pedicel growth phenotype or stomatal phenotype (stomatal index) of *pub30 pub31* (Fig. 3f-i). Therefore, the E3 ligase activity of PUB30/31 is indeed required for the proper pedicel elongation and stomatal development. Based on these findings, we conclude that PUB30 and PUB31 mediate ERECTA ubiquitination both *in vitro* and *in vivo* and regulate the accumulation of ligand-stimulated ERECTA protein.

### Co-receptor of ERECTA, BAK1, interacts with PUB30 and PUB31

EPF/EPFL ligands trigger the active receptor complex formation of ERECTA and its co-receptor BAK1^20^. We thus sought to decipher the regulatory relationships between BAK1 and PUB30/31. First, we asked whether BAK1 could directly interact with PUB30/31. As show in Fig. 4a, the cytosolic domain of BAK1 fused with DNA-binding domain (BD-BAK1_CD) interacts with PUB30/31 (AD-PUB30/31) in the Y2H assays. In addition, the *in vitro* pull-down assay confirmed the interaction between recombinant PUB30/31 (GST-PUB30/31) with BAK1_CD (MBP-BAK1_CD) (Supplementary Fig. 5a, b). To quantitatively characterize the interaction property of BAK1_CD with PUB30/31, we further performed the BLI assays (Fig. 4b, c). Compared to ERECTA_CD (Fig. 2b, c), BAK1_CD exhibited approx. 10 times stronger physical interaction with PUB30 and PUB31, albeit at a micromolar level (Fig. 4b, c). We next performed co-immunoprecipitation (Co-IP) analyses to investigate the *in vivo* association of BAK1 with PUB30 and PUB31 in Arabidopsis. Just like the *in vivo* interaction of ERECTA with PUB30/31 (Fig. 2d, e), BAK1 was weakly detected in the absence of the peptide treatment. Upon EPFL6 peptide incubation, however, BAK1 strongly associated with PUB30 and PUB31 (Figs. 4d, e).

**Fig. 4.**
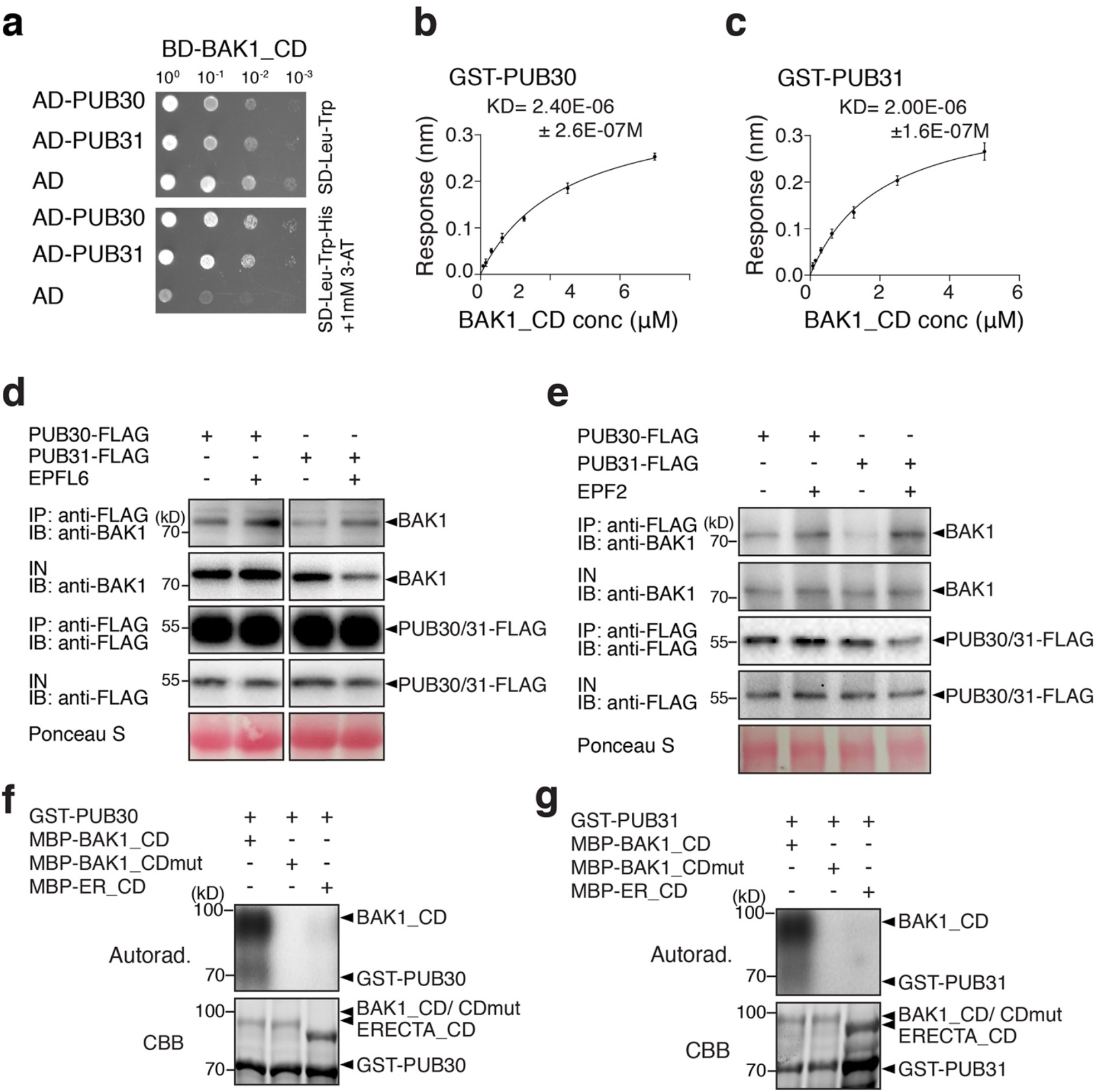
BAK1 interacts with and phosphorylates PUB30 and PUB31. (a) PUB30 and PUB31 interact with cytoplasmic domain of BAK1 (BAK1_CD) in yeast. BAK1_CD were used as bait. PUB30, PUB31, and AD alone were used as prey. Yeast clones were spotted in 10-fold serial dilutions on appropriate selection media. The experiment was repeated independently three times with similar results. (b) A quantitative analysis of interactions between PUB30 and cytoplasmic domain of BAK1 (BAK1_CD) using BLI. *In vitro* binding response curves for recombinantly purified GST–PUB30 and MBP-BAK1_CD at seven different concentrations (78.125 nM, 156.25 nM, 312.5 nM, 625nM, 1250 nM, 2500 nM, and 5000 nM) are shown. Kd values are indicated on the right. Data are mean ± s.d., representative of two independent experiments. (c) A quantitative analysis of interactions between PUB31 and BAK1_CD using BLI. (d) EPFL6 induces the association of PUB30 and PUB31 with BAK1 *in vivo*. After treatment with EPFL6, proteins from *proPUB30::PUB30-FLAG*; *pub30 pub31* and *proPUB31::PUB31-FLAG*; *pub30 pub31* plants were immunoprecipitated with anti-FLAG beads (IP), and the immunoblots (IB) were probed with anti-BAK1 and anti-FLAG antibodies, respectively. (e) EPF2 induces the association of PUB30 and PUB31 with BAK1 *in vivo*. (f) BAK1-CD phosphorylates PUB30 *in vitro*. The phosphorylation of GST-PUB30 was carried out by using MBP-BAK1-CD as the kinase. MBP-BAK1-CDmut was used as a negative control. MBP-ER-CD was also used as kinase for GST-PUB30. Autoradiography (Top) was occupied for phosphorylation detection, and CBB staining (Bottom) was performed to show the protein loading. (g) BAK1-CD phosphorylates PUB31 *in vitro*.

It has been reported that EPF/EPFL signal perceived by ERECTA-BAK1/SERKs is subsequently transduced via BSK1/2 and YODA MAPK cascade^10,47^. To address to which extent PUB30/31 associate with the ERECTA signaling components, we expanded our protein-protein interaction assays. As shown in Supplementary Fig. 6, no interaction of PUB30/31 with BSK1/2 as well as YODA was detected by Y2H. Combined, our results demonstrate that the co-receptor of ERECTA, BAK1, also interacts with PUB30/31 in the EPF/EPFL ligand-stimulated manner and suggest that the regulation of PUB30/31 activity likely occurs at the level of the active receptor complex but not further downstream components.

### BAK1 phosphorylates PUB30 and PUB31

Both ERECTA and BAK1 are functional protein kinases^27,48^ and interact with PUB30/31 upon ligand treatment (Figs. 2, 4). Does ERECTA or BAK1 phosphorylate PUB30/31? To address this question, we first performed *in vitro* kinase assays using purified recombinant epitope-tagged proteins and radioactive ATP (Fig. 4f, g). BAK1_CD (MBP-BAK1_CD) strongly autophosphorylated itself and trans-phosphorylated PUB30/31 (GST-PUB30/31) (Fig. 4f, g). On the other hand, BAK1_CDmut, in which kinase activity is abolished by the substitution of an invariable Lysine to Methionine (K364M), showed no autophosphorylation or phosphorylation of GST-PUB30/31 (Fig. 4f, g). These results suggest that BAK1 phosphorylates PUB30 and PUB31 *in vitro*. We also tested whether ERECTA phosphorylates PUB30/31. However, as reported previously^20^, ERECTA_CD exhibited weak/negligible kinase activity. Consequently, we detected no phosphorylation of PUB30 or PUB31 by ERECT_CD (Fig. 4f, g).

To further identify the exact residue(s) of PUB30/31 phosphorylated by BAK1, we performed liquid chromatography tandem mass spectrometry (LC-MS/MS) analysis after an *in vitro* phosphorylation reaction using MBP-BAK1_CD as kinase and GST-PUB30 as substrate. The threonine 155 (T155) residue, located in the linker domain between the U-box and the ARM repeat domains of PUB30, was identified as a phosphosite (Supplementary Fig. 7a, Supplementary Table 1). The threonine 155 in PUB30 is conserved in PUB31 as threonine 151 (T151) (see Supplementary Fig. 4a). These threonine residues were replaced by alanines (PUB30_T155A_ or PUB31_T151A_) to confirm that they are the major phosphosites. Indeed, GST-PUB30_T155A_ and GST-PUB31_T151A_ were less phosphorylated by BAK1_CD *in vitro* (Supplementary Fig. 7b, c). These results further support that a single amino-acid residues in the linker domain of PUB30/31 are the major *in vitro* phosphosites by BAK1.

### Phosphorylation of PUB30/31 by BAK1 is required for ERECTA-PUB30/31 interaction and ubiquitination of ERECTA

We have shown that EPF/EPFL ligand perception intensifies the association of ERECTA as well as BAK1 with PUB30/31, leading to subsequent ubiquitination and degradation of ERECTA by PUB30/31 (Figs. 2-4). These findings suggest that the ligand-activated ERECTA-BAK1 receptor complex recruits and activates PUB30/31. Because BAK1 directly phosphorylates PUB30/31 *in vitro* (Fig. 4), the important question is whether the BAK1-mediated phosphorylation of PUB30/31 serves as the activation mechanism of these two ubiquitin E3 ligases. To address this question, we first tested whether PUB30_T155_/31_T151_ phosphorylation affects their biochemical activity as E3 ubiquitin ligases. *In vitro* ubiquitination assays were performed using purified recombinant PUB30/31 as well as their phosphomimetic (PUB30_T155D_ and PUB31_T151D_) and phosphonull (PUB30_T155A_ and PUB31_T151A_) versions. As shown in Supplementary Fig. 8, all versions of PUB30 and PUB31 showed similar level of autoubiquitination, suggesting that phosphorylation of PUB30 T155 and PUB31 T151 residues is not required for the E3 ligase activity *per se*.

We sought to further explore if the BAK1-mediated phosphorylation of PUB30 and PUB31 affect their activities in the *in vivo* contexts. To this end, we first examined the effects of PUB30/31 phosphorylation on their interaction with ERECTA. The *in vivo* Co-IP experiments were performed using Arabidopsis protoplasts expressing the epitope-tagged ERECTA and wild-type, phosphomimetic, and phosphonull versions of PUB30/31 (see Methods). The association of ERECTA-HA with PUB30_T155A_-MYC as well as PUB31_T151A_-MYC was markedly reduced compared with the wild-type versions of PUB30/31-MYC (Fig. 5a). In contrast, the phosphomimetic mutant PUB30_T155D_-MYC and PUB31_T151D_-MYC exhibited stronger interactions with ERECTA than the wild-type PUB30 and PUB31, respectively (Fig. 5a). These results indicate that BAK1-mediated phosphorylation of PUB30/31 intensifies their association with ERECTA.

**Fig. 5.**
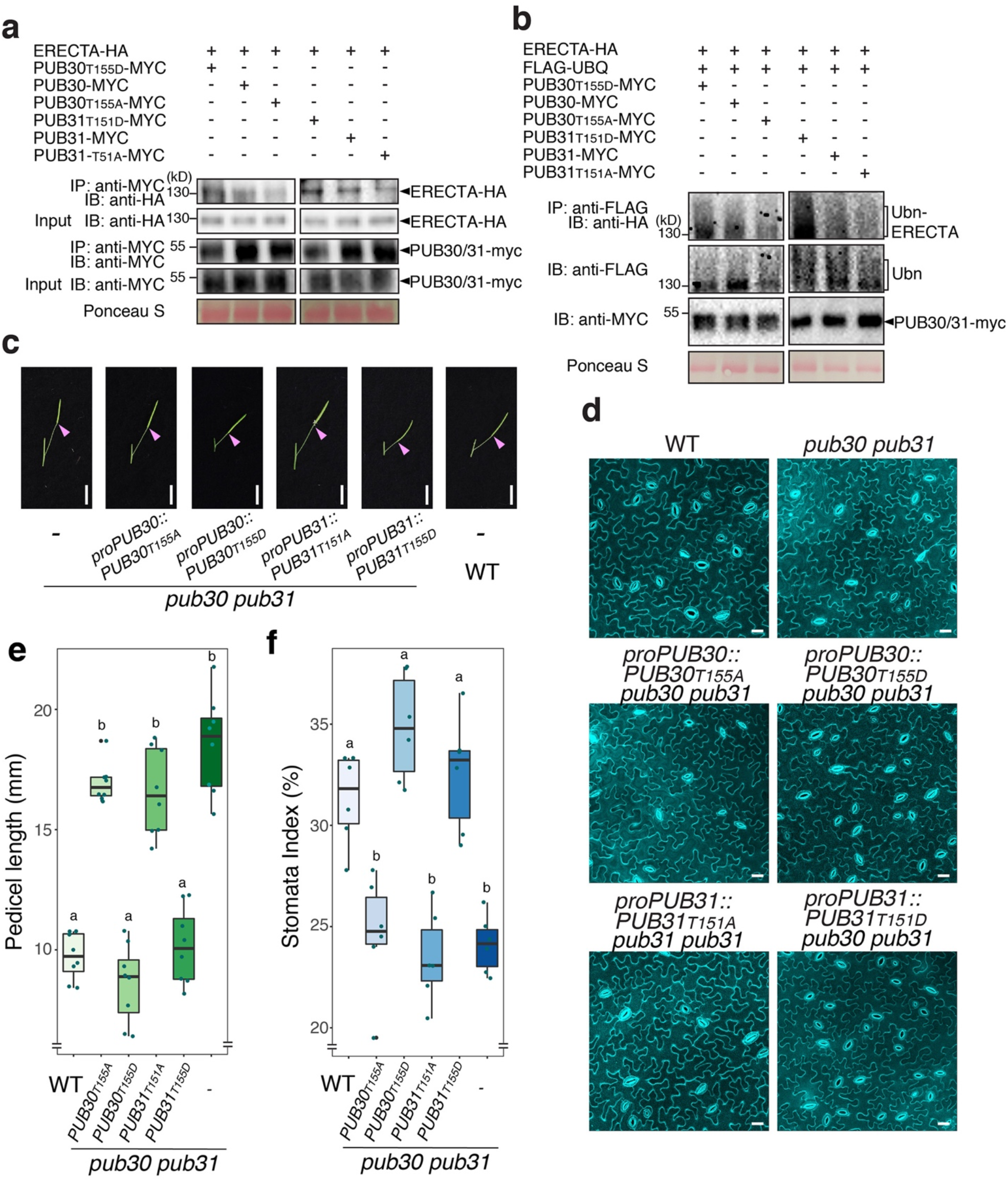
The phosphorylation of PUB30 and PUB31 by BAK1 is essential for the function of PUB30/31. (a) The association of wild-type or various phosphor-mutated PUB30/31 with ERECTA *in vivo*. ERECTA-HA and PUB30 or PUB31 (wild-type or various phosphor-mutated)-MYC plasmids were transfected into protoplast. After incubation for 8h, pretreatment with 2 μM MG132 for 1 h and treatment with 5 μM EPFL6 for 1 h, total proteins were immunoprecipitated with anti-MYC beads (IP), and the immunoblots (IB) were probed with anti-HA and anti-MYC antibodies, respectively. (b) In vivo ERECTA ubiquitination by wild-type or various phosphor-mutated PUB30/31. Arabidopsis protoplasts were co-transfected with ERECTA-HA, FLAG-UBQ, and together with PUB30_T155D_-MYC, PUB30-MYC, PUB30_T155A_-MYC, PUB31_T151D_-MYC, PUB31-MYC, PUB31_T151A_-MYC and incubated for 8 h followed by treatment with 5 μM EPFL6 for 1 h in the presence of 2 μM MG132. After immunoprecipitation using anti-FLAG beads, the ubiquitinated ERECTA was probed with α-HA antibody. The total ubiquitinated proteins were probed by an α-FLAG antibody and PUB30 or PUB31 proteins were probed by an α-MYC antibody. (c) Representative pedicels of mature siliques of *pub30 pub31, proPUB30::PUB30*_*T155A*_; *pub30 pub31, proPUB30::PUB30*T155D; *pub30 pub31, proPUB31::PUB31*_*T151A*_; *pub30 pub31, proPUB31::PUB31*T151D; *pub30 pub31*, and wild type plants. Scale bar: 1 cm (d) Confocal microscopy of 8-d-old abaxial cotyledon epidermis of *pub30 pub31, proPUB30::PUB30*_*T155A*_; *pub30 pub31, proPUB30::PUB30*_T155D_; *pub30 pub31, proPUB31::PUB31*_*T151A*_; *pub30 pub31, proPUB31::*PUB31_T151D_; *pub30 pub31*, and wild type plants. Scale bar: 25 μm (e) Morphometric analysis of pedicel length from each genotype. 6-wk-old mature pedicels (n = 15) were measured. One-way ANOVA followed by Tukey’s HSD test was performed for comparing all other genotypes and classify their phenotypes into two categories (a and b). (f) Quantitative analysis. Stomatal index (SI) of the cotyledon abaxial epidermis from 8-d-old seedlings of respective genotypes. One-way ANOVA followed by Tukey’s HSD test was performed for comparing all other genotypes and classify their phenotypes into two categories (a and b).

Next, we investigated whether the phosphorylation of PUB30 and PUB31 by BAK1 is required for the ubiquitination of ERECTA by PUB30/31. We performed *in vivo* ubiquitination assays in Arabidopsis protoplasts upon EPFL6 peptide application, the condition that triggers polyubiquitination of ERECTA (see Fig. 3e). As evidenced by the reduced ladder-like smear formation, the phosphonull mutants, PUB30_T155A_-MYC and PUB31_T151A_-MYC conferred reduced ubiquitination of ERECTA-HA than the wild-type PUB30-MYC and PUB31-MYC, respectively (Fig. 5b). In contrast, the phosphomimetic mutants, PUB30_T155D_-MYC and PUB31_T151D_-MYC caused increased ubiquitination on ERECTA-HA than the wild-type versions of PUB30/31 (Fig. 5b). Taken together, our results suggest that the BAK1 phosphorylation of PUB30 and PUB31 at T155 and T151 residues, respectively, facilitates the interaction between ERECTA and PUB30/31 and the following ubiquitination of ERECTA by PUB30/31.

### Phosphorylation by BAK1 is required for the function of PUB30 and PUB31

We have shown that BAK1-mediated phosphorylation of PUB30 and PUB31 is critical for subsequent association with and ubiquitination of ERECTA. Do PUB30/31 phosphorylation events affect their *in vivo* functions in plant development? To evaluate the contribution of PUB30 T155 and PUB31 T151 phosphorylation on their biological functions, we introduced the phoshomimetic and phosphonull versions of PUB30/31 driven by their native promoters into the *pub30 pub31* mutant. Transgenic plants expressing *proPUB30::PUB30*_*T155A*_ and *proPUB31::PUB31*_*T151A*_ failed to rescue either pedicel growth or stomatal phenotype of *pub30 pub31* (Fig. 5c-i). In contrast, transgenic plants expressing *proPUB30::PUB30*_*T155D*_ and *proPUB31::PUB31*_*T151D*_ fully rescued the mutant phenotypes, both in the context of pedicel growth and stomatal index (Fig. 5c-f). To exclude the possibility that the absence of phenotypic rescues by the phosphonull mutants of PUB30/31 may be attributed to their reduced protein accumulation, we examined the protein expression levels in these transgenic lines. The PUB30_T155A_/31_T151A_ protein levels were comparable to the phosphomimetic (or wild-type) PUB30/31 (Supplementary Fig. 9). Collectively, our results highlight that the BAK1-mediated phosphorylation of PUB30 and PUB31 is required for their biological functions in regulating inflorescence/pedicel growth and stomatal development.

## Discussion

In this study, we identified a pair of U-box E3 ligases, PUB30 and PUB31, as conserved negative regulators of ERECTA. Our genetic, molecular, and biochemical analyses place PUB30/31 as integral components of the regulatory circuit that fine-tunes the ERECTA signaling outputs (Fig. 6). Upon EPF/EPFL ligand perception by ERECTA, ERECTA forms an active receptor complex with BAK1, which directly phosphorylates PUB30/31. This in turn enhance the interactions between ERECTA and PUB30/31 to trigger ubiquitination and eventual degradation of ERECTA (Fig. 6). Following the activation, the ERECTA-BAK1 complex relays the signal through BSKs, MAPK cascade, and then to the downstream factors to modulate developmental outcomes^23,47,49^. We propose that negative regulation of ERECTA signaling by PUB30/31 ensures robust yet appropriate signaling strengths upon the ligand perception (Fig. 6).

**Fig. 6.**
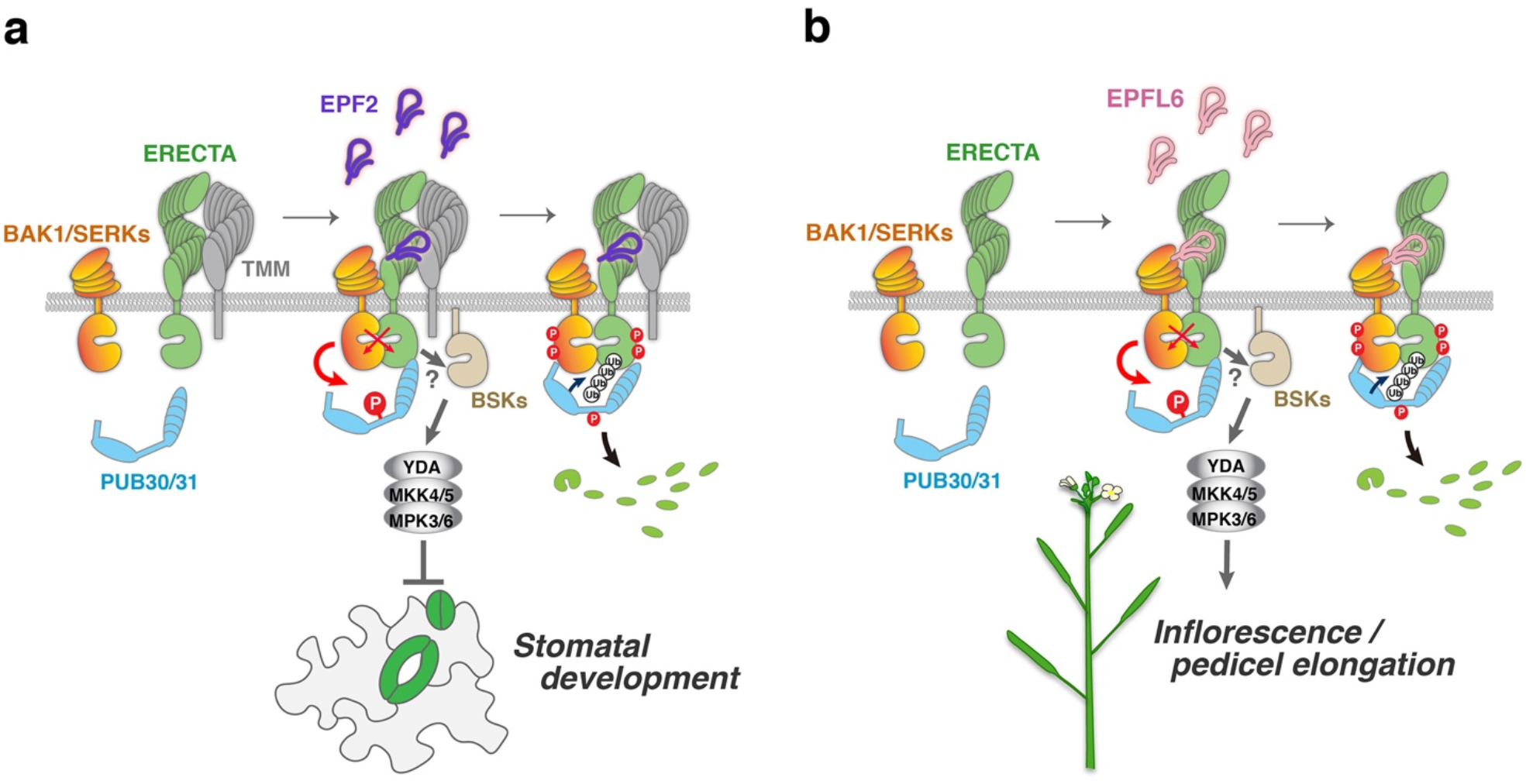
Proposed regulatory mechanisms of ERECTA signal attenuation by PUB30/31 in inflorescence/pedicel growth and stomatal patterning. (a) Regulation of stomatal development. Left: ERECTA (green) and TMM (gray) associate in the absence of ligand. Middle: Upon perception of EPF2 peptide (violet) perception, ERECTA becomes activated, and ERECTA and its co-receptor BAK1/SERKs (orange) undergo transphophorylation events. At the same time, activated BAK1/SERKs phosphorylates PUB30/31 (cyan) at their linker region. The activated ERECTA-BAK1/SERK receptor complex transduces signals most likely (?) via BSK (sand). This leads to the activation of a MAPK cascade composed of YODA-MKK4/5-MPK3/6 and inhibition of stomatal development. Right: The phosphorylated PUB30/31 associate robustly with ERECTA-BAK1/SERK complex, and ubiquitinate ERECTA for eventual degradation. (b) Regulation of inflorescence/pedicel growth. Left: ERECTA (green) and BAK1/SERKs (orange) do not associate strongly in the absence of ligand. Middle: Upon perception of EPFL6 peptide (pink) perception, ERECTA becomes activated, and ERECTA and BAK1/SERKs undergo transphophorylation events. At the same time, activated BAK1/SERKs phosphorylates PUB30/31 (cyan) at their linker region. The activated ERECTA-BAK1/SERK receptor complex transduces signals most likely (?) via BSK (sand). This leads to the activation of a MAPK cascade composed of YODA-MKK4/5-MPK3/6 and promotion of inflorescence and pedicel growth. Right: The phosphorylated PUB30/31 associate robustly with ERECTA-BAK1/SERK complex, and ubiquitinate ERECTA for eventual degradation.

Both ERECTA-mediated inflorescence/pedicel growth and stomatal development are negatively regulated by PUB30 and PUB31 in a largely redundant manner (Figs. 1, S2, 6a, b). The mutations that abolish the E2-binding (PUB30_W97A_ and PUB31_W93A_) as well as the BAK1-mediated phosphosites (PUB30_T155A_ and PUB31_T151A_) uniformly failed to rescue both extremely elongated pedicel and reduced stomatal index phenotypes of the *pub30 pub31* plants (Figs. 3, 5). Thus, whereas each EPF/EPFL peptide ligand elicits a unique developmental response via ERECTA, once the ligands are perceived, subsequent processes of signal activation and attenuation are likely conserved. This appears the case for the EPF2-mediated inhibition of stomatal development (Fig. 6a) and the EPFL6-mediated elongation of inflorescence and pedicels (Fig. 6b). Nonetheless, the ERECTA receptor complex harbors intricate unequal redundancy: ERECTA-LIKE1 (ERL1) and ERL2, two paralogous receptors, synergistically function with ERECTA and form active receptor complexes with SERK1, SERK2, SERK3/BAK1, and SERK4^8,9,20^. It is possible that PUB30/31 exhibit nuanced preferences on these RK family members.

We found that in wild-type plants, ERECTA protein is destabilized most likely via the 26S proteosome pathway and this process relies on PUB30/31, which ubiquitinate the cytoplasmic domain of ERECTA (Fig. 3c, d). This explains the previous finding that a truncated ERECTA protein lacking the entire cytoplasmic domain (ERECTAΔK) is accumulated at high levels, thereby causing dominant-negative effects^38^. Thus, ERECTAΔK is not only unable to transduce signals (owing to the lack of the kinase domain) but also unable be turned over via PUB30/31-mediated ubiquitination. A more recent study has shown that ERL1, after activation by the EPF1 peptide, undergoes rapid internalization via multivesicular bodies/late endosomes to vacuolar degradation^50^. Again, the truncated ERL1 lacking the entire cytoplasmic domain (ERL1ΔK) is stably accumulated at the plasma membrane, irrespective of the ligand perception^50^. It is worth noting that endocytosis of BRI1 relies on the PUB12/13-mediated ubiquitination^31^. Understanding the subcellular dynamics of ERECTA turnover is an interesting area of future research.

Our work broadens the roles of PUB proteins as a regulator of RK signaling and highlights the similarities and differences in their exact mode of actions. We found that ligand-activated BAK1 phosphorylates PUB30/31 at their linker domain (T155 and T151 residues, respectively, Fig. 4). This echoes the idea of phosphorylation as a key activation mechanism of PUB proteins by LRR-RKs and other signaling kinases. For example, MPK3 interacts with and phosphorylates another U-box E3 ligase PUB22^51^; one of which two phosphosites, T88, lies in the linker domain. BRI1 phosphorylates PUB13 at S344, which also falls in the linker domain^31^. This phosphorylation event subsequently facilitates the turn-over of BRI1. Further biophysical and structural analyses may decipher the role of the PUB linker-domain phosphorylation for guiding protein-protein interaction interface with their targets.

The exact steps of phosphorylation-ubiquitination events, on the contrary, are different among the known kinase-PUB signaling modules. For instance, PUB13 ubiquitinates FLS2 upon phosphorylation of PUB13 by BAK1, which in turn strengthens the interaction of PUB13 and FLS2^30^. In contrast, PUB13 ubiquitinates BRI1 in a BAK1-independent manner^31^. Whereas PUB13 marks elicitor-activated FLS2, PUB25/26 specifically target non-activated BOTRYTIS-INDUCED KINASE1 (BIK1), a receptor-like cytoplasmic kinase, for degradation^52,53^. Overall, the regulatory mechanism of the EPF/EPFL-ERECTA-BAK1-PUB30/31 circuit resembles that of flg22-FLS2-BAK1-PUB12/13 but differ from the ubiquitination of BRI1 and BIK1 by PUB12/13 and PUB25/26, respectively.

The inflorescence, pedicel, and stomatal phenotypes of the *pub30 pub31* double mutant plants as well as biochemical analyses indicate that PUB30/31 selectively target ligand-activated ERECTA *in vivo* (Figs 1-3, 5). Surprisingly, PUB30, but not PUB31, has been shown to mediate salt stress tolerance via interacting with and ubiquitinating BRI KINASE INHIBITOR (BKI1)^54^. This raises an important question of what mechanisms govern the target specificities among the PUB proteins. In this regard, it is worth mentioning that PUB12/13, which ubiquitinate the cytoplasmic domains of FLS2 and BRI1 (but not ERECTA)^31^, also interact with and ubiquitinate protein phosphatase 2C (PP2C), ABA-INSENSITIVE1 (ABI1)^55^. The degradation of ABI1 by PUB12/13 promotes ABA signaling and hence drought response^55^. Deciphering the structural basis of association with otherwise unrelated targets will shed light on the versatile roles of PUB proteins in signal transduction pathways underpinning development and environmental responses.

## Methods

### Plant materials and growth conditions

The Arabidopsis accession Columbia (Col) was used as wild type. All plants used in this study are in Col background. T-DNA insertion lines for *PUB30* (SALK_012549) and *PUB31* (SALK_054774) were obtained from Arabidopsis Biological Resource Center. The following mutants and transgenic plant lines were reported previously: *er-105* ^9^; *proERECTA::ERECTA-FLAG* in *er-105*, and *proERECTA::ERECTA-YFP* in *er-105*^*19*^. Arabidopsis seeds were surface sterilized with 30% bleach and grown on half-strength Murashige and Skoog (MS) media containing 1x Gamborg Vitamin (Sigma), 0.75 % Bacto Agar, and 1 % w/v sucrose for 9 days and then transplanted to soil. Plants were grown under long-day conditions (16-h-light /8-h-dark) at 22°C.

### Plasmid construction and generation of transgenic plants

For recombinant protein expression, the following plasmids were generated: pJA51 (MBP-ER_CD), pCLL107 (GST-PUB30), pCLL109 (GST-PUB31), pCLL223 (GST-PUB30_T155A_), pCLL224 (GST-PUB30_T155D_), pCLL225 (GST-PUB31_T151A_), pCLL226 (GST-PUB31_T151D_). To construct GST-PUB30 and GST-PUB31, the coding sequences of PUB30, PUB31 were amplified using Phusion polymerase (Thermo Scientific) and cloned into pGEX-4T-1 using the BamHI and SalI restriction sites. Site-directed mutagenesis was performed using a 2-sided PCR overlap extension followed by assembly into a linearized vector pGEX-4T-1. For Y2H assays, the coding sequences or domain sequences of the genes of interest were fused to either the DNA-binding domain of the pGBKT7 vector or the activation domain of the pGADT7 vector using restriction sites digestion and ligation. The following plasmids were generated: pMM213 (BD-ER_CD), pCLL103 (AD-PUB30), pCLL105 (AD-PUB31), pCLL104 (BD-PUB30), pCLL106 (BD-PUB31), and pCLL143 (BD-BAK1_CD). For complementation assays, the following plasmids were generated: pCLL124 (*proPUB30::PUB30-YFP*), pCLL126 (*proPUB31::PUB31-YFP*), pCLL123 (*proPUB30::PUB30-FLAG*), pCLL125 (*proPUB31::PUB31-FLAG*), pCLL176 (*proPUB30::PUB30*_*W97A*_*-FLAG*), pCLL178 (*proPUB31::PUB31*_*W93A*_*-FLAG*), pCLL235 (*proPUB30::PUB30*_*T155A*_*-FLAG*), pCLL236 (*proPUB30::PUB30*_*T155D*_*-FLAG*), pCLL237 (*proPUB31::PUB31*_*T151A*_*-FLAG*), and pCLL238 (*proPUB31::PUB31*_*T151D*_*-FLAG*). A PCR-based Gateway system was used to generate these constructs. The promoter region (3kb) of *PUB30* and *PUB31* were amplified and subcloned into the pENTR-5’-TOPO cloning vector (Thermo Scientific). The *PUB30* and *PUB31* (WT or mutant) sequences were amplified and subcloned into the pKUT612 cloning vector using restriction enzyme digestion and T4 ligation. Three-way Gateway system^56^ was utilized to generate a series of PUB30 and PUB31 constructs driven by the respective promoters. See Supplemental Tables 2 and 3 for details of plasmid and primer sequence information. Plasmids are transformed into Agrobacterium GV3101/pMP90 and subsequently to Arabidopsis by floral dipping. Over 10 lines were characterized for the phenotypes and reporter gene expressions.

### Real-Time qRT-PCR analysis

RNA extraction, cDNA synthesis, and qRT-PCR were performed as previously described ^57^. For a list of primers, see Supplemental Table 3.

### Histological analysis and microscopy

For histological analysis, mature pedicels were fixed, dehydrated, and embedded into polymethacryl resin Technovit 7100 (Heraeus Kulzer, Wehrheim, Germany) as described previously^38^. Tissue sections were prepared using Leica RM-6145 microtome, and tissue sections were stained with 0.1 % toluidine blue (Sigma) in 0.1 M NaPO_4_ buffer (pH 7.0) and observed under Olympus BX40 light microscope. Confocal microscope images were taken using either Zeiss LSM700 operated by Zen2009 (Zeiss) described previously^19^ or Leica SP5-WLL operated by LAS AF (Leica). Cell peripheries of seedlings were visualized with propidium iodide (Molecular Probes). Images were taken with excitation at 514 nm and emission at 518–600 nm for YFP, and excitation at 619 nm and emission at 642 nm for propidium iodide. For SP-5, a HyD detector was used. The confocal images were false colored, and brightness/contrast were uniformly adjusted using Photoshop 2021 (Adobe). The Z-stack projection images were taken at the interval of 0.99 μm, covering the thickness of the entire cotyledon.

### Quantitative analysis

For analysis of epidermis, abaxial cotyledons from 10-d-old, 6-d-old, or 8-d-old seedlings of respected genotypes were subjected to PI staining and confocal microscopy. The central regions overlying the distal vascular loop were imaged and numbers of epidermal cells, stomata and their cluster size were quantified. Pedicel lengths were measured using ImageJ. Statistical analysis was performed using R ver. 4.1.0 operated under R-Studio ver. 1.4.1717 (https://www.rstudio.com), and graphs were generated using R ggplot2 package.

### Yeast two-hybrid assay

Bait and prey constructs were co-transformed into the yeast strain AH109 using the yeast transformation kit (Frozen-EZ Yeast Transformation II Kit, Zymo Research). The resulting transformants with appropriate positive and negative controls were spotted on SD (−Leu, −Trp) plates to check for growth in the absence of selection. The transformants were then spotted on SD (−Trp, −Leu, −His) selection media containing 1 mM 3-amino-1,2,4-triazole (Sigma, A8056). The positive interactors were then scored based on the stringency of the selection.

### Expression, purification, and refolding of peptides

Recombinant MEPF2 and MEPFL6 peptides were prepared as described previously^19^. Bioactivities of refolded peptides were confirmed as described priviously^19^.

### Co-immunoprecipitation, protein gel electrophoresis and immunoblots

For co-immunoprecipitation assays with seedlings, Arabidopsis transgenic lines expressing different combinations of *proERECTA::ERECTA-YFP, Lti6B-GFP, proPUB30::PUB30-FLAG* and *proPUB31::PUB31-FLAG* were generated. For peptide treatment, Arabidopsis seedlings were grown for five days on ½ MS media plates and then transferred to ddH2O for 24 h. Seedlings were firstly treated with 50 μM MG132 (Sigma) for 3 h. Thereafter, further treatment was performed with Tris-HCl (pH8.0, 50mM) buffer only, MEPF2 (2.5 μM) or MEPFL6 (2.5 μM) at room temperature for another 3 h before being pooled for harvest and then subjected to protein preparation.

The samples were ground in liquid nitrogen and homogenized in the extraction buffer (100 mM Tris-HCl pH 8.8, 150 mM NaCl, 1 mM EDTA, 20% glycerol, 1 mM PMSF, 1x cOmplete protease inhibitor cocktail from Roche, 1x phosphatase inhibitor cocktail 2 and 3 from Sigma). The slurry was centrifuged at 10,000 *g* for 15 min at 4°C. The supernatant was sonicated on ice and then centrifuged at 100,000 *g* for 30 min at 4°C to yield microsomal fractions. The pellet was resuspended in membrane solubilization buffer (100 mM Tris-HCl at pH 7.3, 150 mM NaCl, 1 mM EDTA, 10% glycerol, 1% Triton X-100, 1 mM PMSF, 1x cOmplete protease inhibitor cocktail from Roche, 1x phosphatase inhibitor cocktail 2 and 3 from Sigma) to release membrane proteins. The solution was sonicated on ice and centrifuged again at 100,000 g for 30 min at 4°C. The supernatant was incubated with Protein G-coupled magnetic beads (Dynabeads Protein G, Invitrogen) that captured anti-FLAG (ab205606; Abcam) antibody at 4°C for 2h with gentle agitation. Then, the beads were washed four times with 500 μl of phosphate buffer (pH 7.4) and precipitated proteins were eluted with 4x SDS sample buffer at 95 °C for 5 minutes. Either total membrane or immunoprecipitated proteins were separated on a SDS-PAGE gel and transferred to PDVF membrane (Millipore) for immunoblot analysis using monoclonal anti-GFP (33-2600, 1:1,000, Thermo Fisher Scientific), anti-FLAG (F-3165, 1:5,000, Sigma), and anti-BAK1 (AS12 1858, 1:5,000, Agrisera) as primary antibodies. As secondary antibodies, sheep anti-mouse IgG horseradish peroxidase-linked antibody (NA931, GE Healthcare) and goat anti-rabbit IgG (whole molecule)–peroxidase antibody (A6154, Sigma) were used at a dilution of 1:50,000 and 1:6,000, respectively. The protein blots were visualized using Chemi-luminescence assay kit (34095, Thermo Scientific).

For co-immunoprecipitation assays with Arabidopsis protoplasts, protoplasts were transfected with HA-tagged ERECTA (ER-HA) and MYC-tagged PUB30 or PUB31 (wild-type or various phosphor-mutated) and incubated for 8 h. Then, protoplasts were pretreated with 2 μM MG132 (Sigma) for 1 h, followed by treatment with 5 μM EPFL6 for 1 h. The total proteins were isolated with extraction buffer (50 mM Tris·HCl, pH 7.5, 150 mM NaCl, 1 mM EDTA, 20% glycerol, 1% Triton X-100, and 1x cOmplete protease inhibitor cocktail from Roche, 1x phosphatase inhibitor cocktail 2 and 3 from Sigma). The supernatant was incubated with Protein G-coupled magnetic beads (Dynabeads Protein G, Invitrogen) that captured anti-MYC (ab9106; Abcam) antibody at 4°C for 2h with gentle agitation. Then, the beads were washed three times with 500 μl of wash buffer (50 mM Tris·HCl, pH 7.5, 150 mM NaCl, 1 mM EDTA, 20% glycerol, 0.2% Triton X-100, and 1x cOmplete protease inhibitor cocktail from Roche, 1x phosphatase inhibitor cocktail 2 and 3 from Sigma) and precipitated proteins were eluted with 4x SDS sample buffer at 95 °C for 5 minutes. Either total or immunoprecipitated proteins were separated on a SDS-PAGE gel and transferred to PDVF membrane (Millipore) for immunoblot analysis using anti-HA (ab18181, 1:1,000, Abcam), and anti-MYC (ab32, 1:1,000, Abcam) as primary antibodies. As secondary antibody, goat anti-mouse IgG H&L (HRP) (ab205719, Abcam) was used at a dilution of 1:10,000. The protein blots were visualized using Chemi-luminescence assay kit (34095, Thermo Scientific).

### *In vitro* pull-down assay

For pull-down assay of PUB30/31 and ER_CD, approx. 15 μg GST-PUB30 or GST-PUB31 or GST proteins were incubated with about 15 μg MBP-ERECTA_CD protein in 900 μl pull-down buffer (10% glycerol, 1% Triton X-100, 1.5mM MgCl2, 150mM NaCl, 1mM EGTA, 50mM HEPES, pH 7.5, 1 mM PMSF and 1× cOmplete protease inhibitor cocktail from Roche). 30 μl GST beads (Glutathione Sepharose 4 Fast Flow, 17-5132-01, Cytiva) were incubated with each reaction mixture with gentle shaking at 4°C for about 1 h. For pull-down assay of PUB30/31 and BAK1_CD, approx. 15 μg GST-PUB30 or GST-PUB31 proteins were incubated with about 15 μg MBP-BAK1_CD or MBP proteins in 900 μl pull-down buffer. 30 μl MBP beads (Amylose Resin, E8021S, New England Biolabs) were incubated with each reaction mixture with gentle shaking at 4°C for about 1 h.

After reaction, beads were washed three times and heated for 5 min in a 95 °C metal bath. The immunoprecipitated proteins were separated by SDS-PAGE electrophoresis and detected by anti-GST (A00865-200, 1:5,000, Genscript) and anti-MBP (E8032, 1:10,000, New England Biolabs) antibodies, respectively. As secondary antibody, sheep anti-mouse IgG horseradish peroxidase-linked antibody (NA931, GE Healthcare) was used at a dilution of 1:50,000. The protein blots were visualized using Chemi-luminescence assay kit (34095, Thermo Scientific).

### *In vitro* and *in vivo* ubiquitination assays

The *in vitro* ubiquitination reactions contain 1 μg each of MBP-ERECTA_CD, HIS6-E1 (AtUBA1), HIS6-E2 (AtUBC8), HIS6-ubiquitin (Boston Biochem), and GST-PUB30 or GST-PUB31 in the ubiquitination reaction buffer (0.1 M Tris-HCl, pH 7.5, 25 mM MgCl2, 2.5 mM DTT, and 10 mM ATP; final volume 30 μl). The reactions were incubated at 30 °C for 3h and then stopped by adding SDS sample loading buffer and boiled at 95 °C for 5 min. The samples were then separated by 8% SDS-PAGE, and the ubiquitinated ERECTA_CD were detected by IB analysis with anti-MBP (E8032, 1:10,000, New England Biolabs) as primary antibody, whereas the autoubiquitination was detected by IB analysis with anti-GST (A00865-200, 1:5,000, Genscript) as primary antibody. As secondary antibody, sheep anti-mouse IgG horseradish peroxidase-linked antibody (NA931, GE Healthcare) was used at a dilution of 1:50,000. The protein blots were visualized using Chemi-luminescence assay kit (34095, Thermo Scientific).

For *in vivo* ubiquitination assays, Arabidopsis protoplasts were co-transfected with FLAG-tagged ubiquitin (FLAG-UBQ), HA-tagged ERECTA (ER-HA) and together with a control vector or MYC-tagged PUB30 or PUB31 (wild-type or various phosphor-mutated) and incubated for 8 h followed by treatment with 5 μM EPFL6 for 1 h in the presence of 2 μM MG132 (Sigma). The ubiquitinated ERECTA was detected with an α-HA (ab18181, 1:1,000, Abcam) IB after IP with α-FLAG (ab205606, Abcam) antibody. The total ubiquitinated proteins were detected by anti-FLAG (F-3165, 1:5,000, Sigma) and α-MYC (ab32, 1:1,000, Abcam) as primary antibodies. As secondary antibody, sheep anti-mouse IgG horseradish peroxidase-linked antibody (NA931, GE Healthcare) was used at a dilution of 1:50,000. The protein blots were visualized using Chemi-luminescence assay kit (34095, Thermo Scientific).

### *In vitro* kinase assay

Kinase assays were conducted in 30 μl reactions containing 20 mM Tris-HCl (pH7.5), 5 mM EGTA, 1 mM DTT, 100 mM NaCl, 10 mM MgCl_2_, 100 μM ATP [γ-32P] mix (5μCi of ATP, ATP, [γ-32P]-3000 Ci/mmol 10m Ci/ml EasyTide, 100 μCi) and 10 μg of substrate proteins and 1 μg of kinases. The reactions were incubated at 30 ºC for 30 minutes and stopped with the SDS sample buffer. After the SDS-PAGE, gels were dried, subsequently exposed to a GE Multipurpose Standard Screen (63-0034-87) for 18 hours and imaged using a GE Typhoon FLA 9000 Gel imager.

### Mass-spectrometry and identification of the phosphosites

For *in vitro* phosphorylation reaction, 1 μg MBP-BAK1_CD protein was incubated with 10 μg GST-PUB30 in 30 μl kinase reaction buffer at 30 °C for 3 h (with gentle shaking). After the reaction, the SDS loading buffer was used to stop the kinase reactions. Samples were separated by 10% SDS– PAGE. The gels were fixed 60 minutes in 50% methanol + 7% acetic acid, rinsed thoroughly with Milli Q water and stained with GelCode Blue Stain Reagent (Thermo Fisher cat # 24590). Target bands for GST-PUB30 are cut off from the electrophoresis gel and digested using chymotrypsin (Sigma) at 37 °C overnight. To analyze the chymotryptic peptides, nano-flow reverse phase liquid chromatography and tandem MS was performed using Q Exactive Hybrid Quadrupole-Orbitrap Mass Spectrometer (ThermoFisher Scientific) as described previously^58^. Subsequently, the peptide identification was performed by searching the *Arabidopsis thaliana* reference genome (downloaded from https://www.uniprot.org) using SEQUEST (ThermoFisher Scientific) search engine. The parameter of dynamic modifications with phosphorylation filter was added for the identification of phosphopeptides. The peptide spectrum match details with phosphorylated residues were manually validated and annotated in Thermo Proteome Discoverer 2.4.

### Proteasome inhibitor treatment and immunoblot Assays

The *proERECTA::ERECTA-FLAG er-105* and *proERECTA::ERECTA-FLAG pub30 pub31 er-105* seedlings were grown vertically at 22°C on half-strength Murashige and Skoog (MS) medium for 5 d and then transferred to liquid half-strength MS medium with or without 20μM MG132 (Sigma) for 48 h. Total protein extracts were separated on a 10% SDS-polyacrylamide gel and detected by immunoblot analysis with anti-FLAG (F-3165, 1:5,000, Sigma) and anti-actin (ab230169, 1:2,000, Abcam) as primary antibodies. As secondary antibody, sheep anti-mouse (NA931, GE Healthcare) was used at a dilution of 1:50,000. The protein blots were visualized using Chemi-luminescence assay kit (34095, Thermo Scientific).

### Biolayer interferometry (BLI)

The binding affinities of the ERECTA_CD with GST-tagged PUB30 and PUB31 were measured using the Octet Red96 system (ForteBio, Pall Life Sciences) following the manufacturer’s protocols. The optical probes coated with anti-GST were first loaded with 2000 nM GST-PUB30 or PUB31 before kinetic binding analyses. The experiment was performed in 96-well plates maintained at 30 °C. Each well was loaded with 200 μl reaction volume, and the binding buffer used in these experiments contained 1× PBS supplemented with 0.02 % Tween 20. The concentrations of the ERECTA_CD as the analyte in the binding buffer were 20000 nM, 10000 nM, 5000 nM, 2500 nM, 1250 nM, 625 nM, and 312.5 nM. Similarly, for the binding of BAK1-CD with GST-PUB30/31, the optical probes coated with anti-GST were first loaded with 1000 nM GST-PUB30 or PUB31 before kinetic binding analyses. The concentrations of the BAK1_CD as the analyte in the binding buffer were 5000 nM, 2500 nM, 1250 nM, 625 nM, 312.5 nM, 156.3 nM, and 78.2 nM. All preformed complexes remained stable as suggested by the constant signal during the washing step after loading. There was no binding of the analytes to the unloaded probes as shown by the control wells. Binding kinetics to all seven concentrations of the analytes were measured simultaneously using default parameters on the instrument. The data were analyzed using the Octet data analysis software. The association and dissociation curves were fit with the 1:1 homogeneous ligand model. The kobs (observed rate constant) values were used to calculate *K*_d_, with steady-state analysis of the direct binding.

## Acknowledgements

We thank Dr. Keiko Kuwata and Dr. Pengfei Bai for assistance in mass-spec analysis, Dr. Krishna Sepuru for training of BLI, Dr. Julian Avila and Dr. Michal Maes for ERECTA expression plasmids, Prof. Libo Shan, and Prof. Martin Bayer for sharing published materials, Dr. Mireia Panicot Anglada for initial screens of T-DNA lines, and members of the Torii Lab for discussion. This work was initially supported by the US National Science Foundation (IOS-0744892) and then by the HHMI and UT Austin, Johnson & Johnson Centennial Chair to K.U.T. J.M.W. was supported by Mary Gates Undergraduate Research Fellowship from the University of Washington.

## Author contributions

Project conception: K.U.T., K.S.; Conceptualization, L.-L.C., K.U.T.; Experimental Design, L.-L.C., K.U.T.; Performance of experiments, L.-L.C., A.M.C., S.M.B., J.M.W., K.U.T.; (genetic/phenotypic analyses, L-L.C., A.M.C., S.M.B., K.S., J.M.W., K.U.T.; Biochemical/biophysical/molecular experiments, A.M.C., L-L.C); Data analysis, L.-L.C., K.U.T.; Visualization, L.-L.C., K.U.T.; Writing, L.-L.C., K.U.T.; All authors contributed to the editing of the manuscript; Project Administration, K.U.T.; Funding Acquisition, K.U.T.

